# Optimized fragmentation improves the identification of peptides cross-linked using MS-cleavable reagents

**DOI:** 10.1101/476051

**Authors:** Christian E. Stieger, Philipp Doppler, Karl Mechtler

## Abstract

Cross-linking mass spectrometry (XLMS) is becoming increasingly popular, and current advances are widening the applicability of the technique so that it can be utilized by non-specialist laboratories. Specifically, the use of novel mass spectrometry-cleavable (MS-cleavable) reagents dramatically reduces complexity of the data by providing i) characteristic reporter ions and ii) the mass of the individual peptides, rather than that of the cross-linked moiety. However, optimum acquisition strategies to obtain the best quality data for such cross-linkers with higher energy C-trap dissociation (HCD) alone is yet to be achieved. Therefore, we have carefully investigated and optimized MS parameters to facilitate the identification of disuccinimidyl sulfoxide (DSSO)- based cross-links on HCD-equipped mass spectrometers. From the comparison of 9 different fragmentation energies we chose several stepped-HCD fragmentation methods that were evaluated on a variety of cross-linked proteins. The optimal stepped-HCD-method was then directly compared with previously described methods using an Orbitrap Fusion™ Lumos™ Tribrid^TM^ instrument using a high-complexity sample. The final results indicate that our stepped-HCD method is able to identify more cross-links than other methods, mitigating the need for multistage MS (MS^n^) enabled instrumentation and alternative dissociation techniques.

---

Cross-linking mass spectrometry (XLMS) is a rapidly growing field of research at the interface of proteomics and structural biology.^1–5^ Typically, N-hydroxysuccinimide (NHS)-based functionalities that link primary amines (lysine, protein N-terminus) and hydroxyl groups (serine, threonine and tyrosine) are used as reactive groups to form covalent bonds between residues that are in close spatial proximity. After reacting proteins, protein complexes or even whole cells with one of these reagents, the sample is digested enzymatically. This results in a complex mixture containing linear peptides, mono- or dead-end-links (peptides that reacted with one end of the cross-linker while the other reactive group is hydrolyzed), intra-peptide cross-links (peptides containing two linkable amino acids without enzymatic cleavage site in-between) and inter-peptide cross-links (cross-links that link two separate peptides). This mixture is then analyzed via high performance liquid chromatography (LC) -tandem mass spectrometry (MS/MS or MS^2^) to identify the cross-linked species. Since cross-linked peptides are formed sub-stoichiometrically, mass spectrometers offering high sensitivity and high scan rates are required for a comprehensive identification.

As well as their detectability, the estimation of the false discovery rate (FDR) for cross-linked peptides is also more challenging compared to linear peptides.^6,7^ Over the last decade, several MS-cleavable cross-linker have been developed, that facilitate data analysis and diminish the possibility of false positives.^8–11^ The two most commonly used and commercially available MS-cleavable cross-linker, disuccinimidyl sulfoxide (DSSO, Fig. 1) and disuccinimidyl dibutyric urea (DSBU or BuUrBu, Fig. 1) have been extensively investigated and are increase the reliability of XLMS results. Both contain chemical groups that cleave upon collisional activation. DSBU is the diamide of carbonic acid and aminobutanoic acid. These amide bonds have a stability comparable to that of the amide bonds in the peptide backbone. Therefore, higher energy C-trap dissociation (HCD), a beam-type collision-induced dissociation method, is the fragmentation method of choice and is frequently applied in the measurement of DSBU cross-linked peptides.^12,13^ In contrast, the C-S bonds adjacent to the sulfoxide group of DSSO is weaker than the peptide backbone and can be selectively cleaved upon collisional induced dissociation (CID).

**Figure 1.**
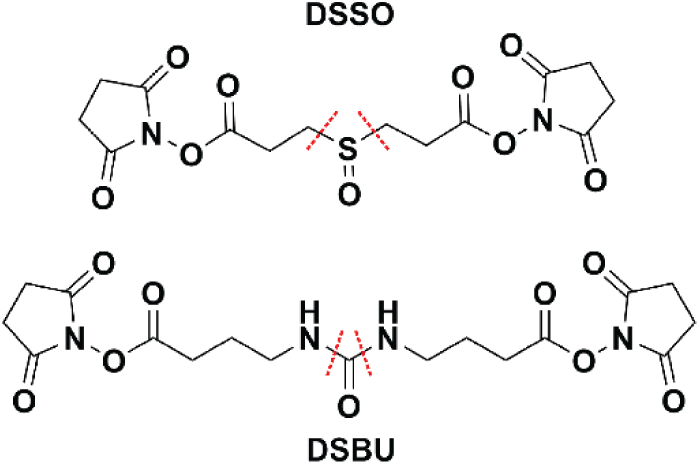
Illustration of disuccinimidyl sulfoxide (DSSO) and disuccinimidyl dibutyric urea (DSBU). Red dotted lines indicate collisional induced dissociation associated cleavage sites.

To obtain satisfying sequence coverage, required for unambiguous cross-link identification, a second MS/MS-scan with a complementary fragmentation method such as electron-transfer dissociation (ETD), or separate sequencing of the arising reporter doublets in 3^rd^ stage mass spectrometry (MS^3^) is needed.^14,15^ The disadvantage of sequential MS/MS scans or multistage MS (MS^n^) methods are reduced scan rate and lower sensitivity. Moreover, advanced instruments like Orbitrap Fusion or Orbitrap Fusion Lumos are required for optimal performance. Previous experiments have already described HCD to yield the highest number of identified cross-links for non-cleavable cross-linking reagents.^16,17^ These studies impressively demonstrated, that optimal fragmentation is crucial for cross-link analysis and allows the identification of up to 4 times more cross-linked peptide pairs. Moreover, they highlight HCD to be the fragmentation strategy of choice for most cross-linked species.

A recent publication by Smith et al.^18^ gives a detailed comparison on different fragmentation methods, and data analysis tools for peptides cross-linked with both DSSO and DSBU. This comparison also includes two different fragmentation approaches namely, stepped-HCD and sequential CID-ETD fragmentation. However, this study did not investigate the collision energy dependency on the fragmentation of DSSO-cross-linked peptides and optimization of the stepped-HCD acquisition strategy was not performed. Moreover, in this study two particular proteins have been used, that yielded a limited number of cross-links. In the study presented herein, we have elucidated the influence of different normalized collision energies (NCEs) on the HCD-fragmentation behavior of DSSO cross-linked peptides. Furthermore, we optimized the fragmentation energy to obtain the best possible results on HCD-cell equipped mass spectrometers.

Optimization was carried out on cross-linked peptides derived from five different proteins rabbit aldolase, bovine serum albumin (BSA), equine myoglobin, *S. pyogenes* Cas9 and human transferrin. Cross-link precursors were identified by using the previously published sequential CID-ETD acquisition method and XLinkX 2.2 for database search.^15^ On the basis of fragmentation behavior, search engine scores and identified cross-linked species, different stepped collision energies were proposed and compared to each other. Finally, the optimal performing stepped collision energy was compared to published acquisition strategies for DSSO^14,15^ using two commercially available systems; BSA which served as a model system for a single protein and the 70S *E. coli* ribosome that served as a model system for a more complex sample.

## MATERIALS AND METHODS

### Chemicals and Reagents

Aldolase (rabbit), alcohol dehydrogenase (ADH, yeast), bovine serum albumin (BSA), conalbumin (chicken), myoglobin (equine), ovalbumin (chicken) and transferrin (human) were purchased from Sigma-Aldrich (St. Louis, MO, USA), *E. coli* 70S ribosome from New England Biolabs (Ipswich, MA, USA) and DSSO from Thermo Fisher Scientific (Rockford, IL, USA). Cas9 (*S. pyogenes*) with a fused Halo-tag was expressed and purified as described by Deng et al.^19^

### Cross-Linking and Digestion

Prior to cross-linking Cas9 was buffer exchanged to XL-buffer (50 mM HEPES, 50 mM KCl, pH 7.5) using Micro Bio-Spin™ 6 columns (Bio-Rad, Hertfordshire UK). The other proteins were dissolved in XL-buffer (the ribosome sample contained additionally 10 mM MgAc_2_). All proteins were cross-linked separately at a concentration of 1 µg/µl and a final DSSO concentration of 500 µM. The reaction was carried out for 1 h at room-temperature, before it was quenched with 50 mM Tris-HCL, pH 7.5. Samples were reduced with dithiothreitol (DTT, 10 mM, 30 min, 60 °C) and alkylated with iodoacetamide (IAA, 15 mM, 30 min at room temperature in the dark). Alkylation was stopped by the addition of 5 mM DTT and proteins were digested with trypsin overnight (protein/enzyme 30:1, 37 °C). Digestion was stopped by the addition of trifluoracetic acid to a final concentration of 1% (v/v, pH <2) and samples were stored at −80 °C. The sample for stepped-HCD comparison was obtained by mixing all digestsin an equimolar ratio.

For the sequential digest, tryptic peptides were desalted using self-made C-18 Stage Tips^20^, dissolved in XL-buffer and digested with *S. aureus* Protease V8 (GluC) for 4 hours (protein/enzyme 30:1, 37 °C).

### Size-exclusion-chromatography (SEC) enrichment

The ribosome sample was enriched for XLs prior to LC-MS/MS analysis using SEC. 25 µg of the digest were separated on a TSKgel SuperSW2000 column (300 mm × 4.5 mm × 4 μm, Tosoh Bioscience). (Figure S8) The three high mass fractions were subsequently measured via LC-MS/MS.

### Reversed-Phase HPLC

Digested peptides were separated using a Dionex UltiMate 3000 HPLC RSLC nanosystem prior to MS analysis. The HPLC was interfaced with the mass spectrometer via a Nanospray Flex™ ion source. For sample concentrating, washing and desalting, the peptides were trapped on an Acclaim PepMap C-18 precolumn (0.3×5mm, Thermo Fisher Scientific), using a flowrate of 25 µl/min and 100% buffer A (99.9% H2O, 0.1% TFA). The separation was performed on an Acclaim PepMap C-18 column (50 cm x 75 µm, 2 µm particles, 100 Ä pore size, Thermo Fisher Scientific) applying a flowrate of 230 nl/min. For separation, a solvent gradient ranging from 2-35% buffer B (80% ACN, 19.92% H2O, 0.08% TFA) was applied. The applied gradient varied from 60-180 min, depending on the sample complexity.

### Mass Spectrometry

All measurements were performed on an Orbitrap Fusion™ Lumos™ Tribrid™ (Thermo Fisher Scientific) mass spectrometer.

### Data Dependent Acquisition Methods

For DSSO-XL identification, digested proteins were analyzed using the CID-ETD acquisition method described by Liu et al.^14^ Full scans were recorded at resolution 60000 and a scan range from 375-1500 *m/z* (AGC 4e5, max injection time 50 ms). MS/MS scans were recorded at 30000 resolution (AGC 5e4, max injection time 100 ms for CID and 120 ms ETD, isolation width 1.6 *m/z*). Singly and doubly charged ions were excluded from fragmentation since cross-linked peptides tend to occur at a charge state of 3+ or above.^21^ CID fragmentation energy was set to 25 % NCE and for ETD calibrated charge dependent ETD parameters were used. In CID-ETD acquisition, two subsequent fragmentation events using the complementary fragmentation strategies were triggered. All precursors that have been selected for fragmentation were excluded from fragmentation for 30 s.

For the MS^n^ acquisition strategy (from now on called MS^2^-MS^3^) the same settings as described above were used but only precursor with charge state 4-8+ were selected for MS/MS. The two most abundant reporter doublets from MS/MS scans (charge state 2-6, Δ-mass 31.9721 Da, ±30 ppm) were selected for MS^n^. MS^3^ scans were recorded in the ion trap operated in rapid mode with a maximum fill time of 150 ms (isolation width 2.0 *m/z*). Fragmentation was carried out using HCD with 30 % NCE.

For stepped-HCD the settings described above were used with one adaptation. Ions for MS/MS were collected for a maximum of 150 ms. Selected precursors were fragmented applying a collision energy of 27±6 % NCE.

### NCE optimization

An inclusion list was generated that included all cross-links identified with CID-ETD using the Proteome Discoverer 2.2 output. Full scans were recorded at a resolution of 120000 ranging from 400-1600 *m/z* (AGC 2e5, 50 ms max. injection time). Only precursors from the inclusion list (10 min retention time window, matching charge and *m/z* [± 10 ppm]) were selected for fragmentation. MS/MS spectra were recorded at 30000 resolution (AGC 1e5, max. injection time 150 ms, isolation width 1.4 *m/z*). Each selected precursor was fragmented consecutively with 9 different NCEs and subsequently excluded from fragmentation for 30 s. The chronological order of fragmentation energies was randomly shuffled between the three injection replicates.

To identify the optimal stepped-HCD method for analysis of DSSO-cross-linked peptides, a mixture of 5 proteins was analyzed in triplicate using three different stepped NCEs. To allow an unbiased comparison, and to compensate for chromatographic variations only precursors from an inclusion list (combined inclusion list from separate proteins see “Effect of NCE on Cross-Link identification”) were selected for fragmentation. For each precursor 3 subsequent fragmentation events using different stepped-HCD methods were triggered. Measurements were carried out in triplicate with shuffled fragmentation order.

### Data Analysis

Thermo .raw files were imported into Proteome Discoverer 2.2 and analyzed with XLinkX (version 2.2 or 2.3) using the following settings: Cross-Linker: DSSO (+158.00376 Da, reactivity towards lysine and protein N-terminus for initial identification and NCE optimization, for method comparison serine, threonine and tyrosine were additionally included); cross-linker fragments: alkene (+54.01056 Da), unsaturated thiol (+85.98264 Da), sulfenic acid (+103.9932 Da); cross-link doublets: alkene/unsaturated thiol (Δ-mass 31.96704 Da) or alkene/sulfenic acid (Δ-mass 49.98264 Da); MS1 accuracy: 10 ppm; MS2 accuracy: 20 ppm; MS3 accuracy: 0.5 Da; used enzyme: trypsin; max. missed cleavages: 4; minimum peptide length: 5; max. modifications: 4; peptide mass: 300-7000 Da; static modifications: carbamidomethylation (cysteine, +57.021 Da); dynamic modifications: oxidation (methionine, +15.995 Da). For the database search the false discovery rate (FDR) was set to 1%. To reduce the number of false positives, cross-links identified with XLinkX were filtered for an identification score ≥20 as suggested by Thermo Fisher and additionally for an identification delta score (Δ-score) ≥20. CID-ETD and MS2-MS3 runs were analyzed with the MS2_MS2 or MS2_MS3 workflow provided in Proteome Discoverer 2.2. Detailed analysis parameters using MeroX are described in the supplementary information.

The FASTA files for database search contained the used model proteins and all identified proteins contained in the ribosome sample, respectively.

## RESULTS

### Effect of NCE on XL identification

We first sought to investigate the identification rate of DSSO cross-linked peptides with respect to the NCE employed during HCD activation. Therefore we adopted the workflow described by Kolbowski et al.^16^ (Figure 2). To achieve optimal reproducibility, an inclusion list of previously identified cross-link-precursors was generated for each protein used in the study (Aldolase, BSA, Cas9, Myoglobin, Transferrin). Detection of an XL-precursor triggered nine consecutive fragmentation events utilizing different NCEs, ranging from 15-39% (Figure 2). Samples were measured in triplicate, with randomly shuffled collision energy order. The recorded dataset was analyzed using XLinkX 2.2^15^ and all runs were searched against a database containing the five investigated proteins.

**Figure 2.**
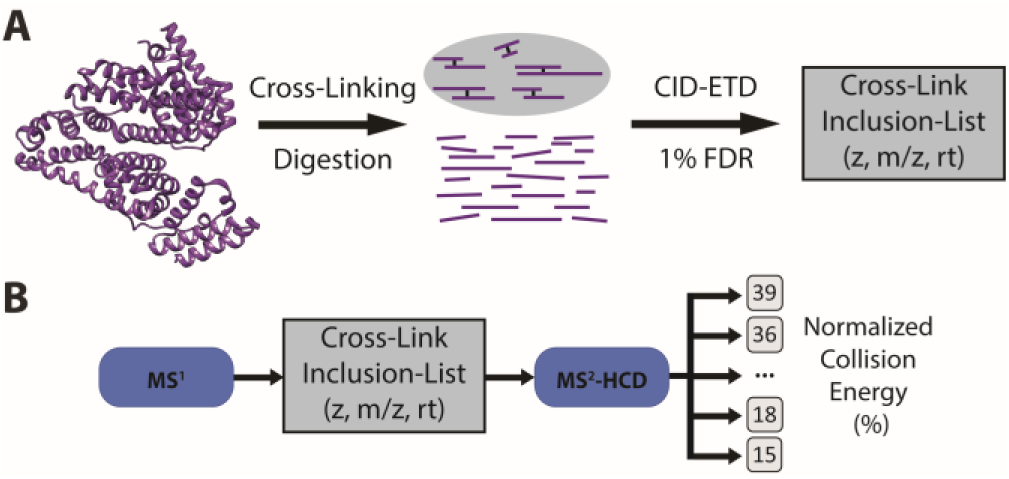
Illustration of the used workflow, adopted from Kolbowski et al.^16^. Proteins were cross-linked separately and analyzed one by one on an Orbitrap Fusion Lumos with the already published sequential CID-ETD method. (A) An inclusion list was generated and used for a subsequent targeted analysis of all identified cross-linked peptides using HCD with different normalized collision energies (B)

On average, NCEs of 21 % and 24 % could identify the highest number of cross-linked sites (e.g. unique linked amino acids; 293/289) when applying a 1% FDR. (Figure 3) Employment of higher NCEs (>27 %) leads to a significantly lower cross-link identification, with 39 % NCE identifying over 70 % less than the two best collision energies (21% and 24%). The score provided by XLinkX reaches its maximum between 27 % and 30 % NCE. (Figure 3A) In addition to XLinkX, the dataset was also analyzed with MeroX 1.6.6.^13^ These algorithms gave similar results, with a slightly shifted ID-maximum at 24 % NCE. (Figure S1).

**Figure 3.**
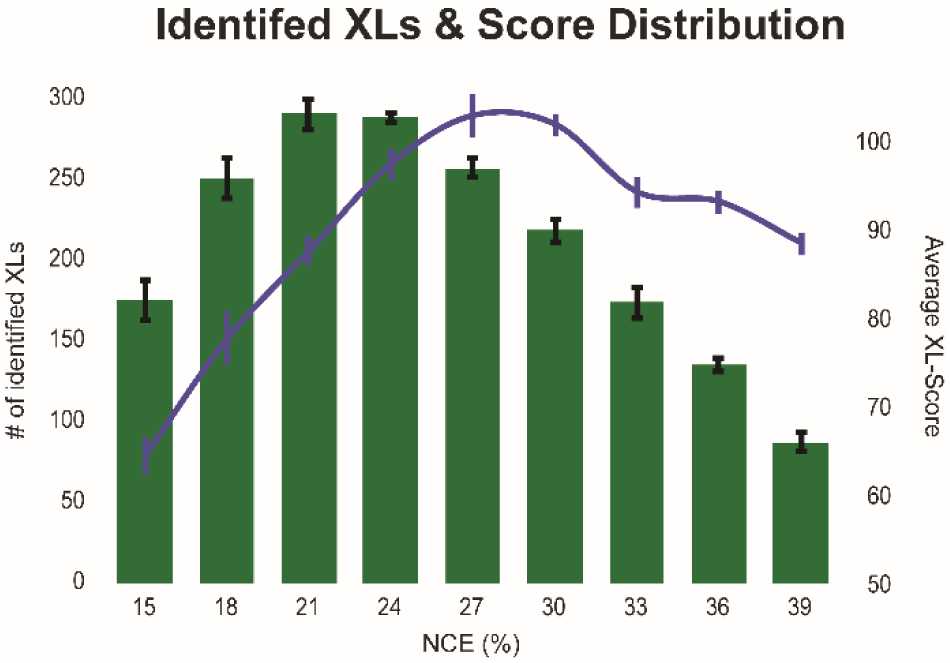
number of identified cross-links (green) and their average score (blue) according to different the normalized collision energies. (n=3, Error bars represent the 0.95 confidence interval [CI])

Since scoring is highly dependent on the scoring-function implemented in the search engine, we aimed for a more independent measure to compare the different fragmentation energies. Hence, we compared the sequence coverage obtained with different NCEs. The data was analyzed with XLinkX 2.3 in PD 2.3 (beta) and the reported sequence coverage for the identified XLs was used for comparing different NCEs. An overall comparison of the sequence coverage already shows that higher NCEs provide higher sequence coverage Figure 4A).

**Figure 4.**
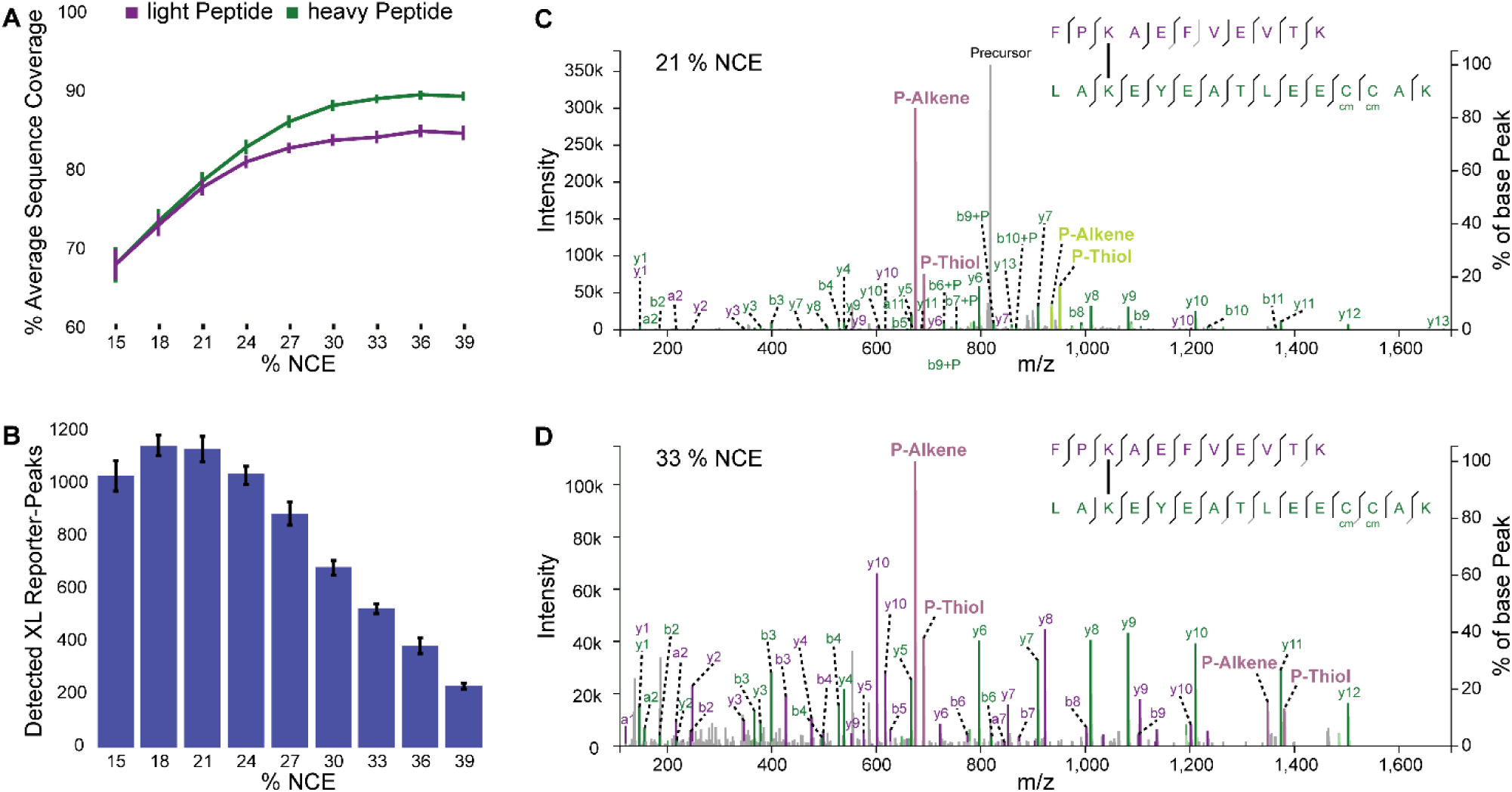
Average Sequence coverage of the identified cross-links at different normalized collision energies. (A) Spectra that have been identified to contain at least two different cross-link reporter doublets as a function of the used normalized collision energy. (B) (n=3, Error bars represent the 0.95 CI) Two MS/MS-spectra triggered from the same precursor using a rather low (C, 21%) and a more elevated (D, 33%) normalized collision energy. Spectra were annotated with the help of xiSPEC (https://spectrumviewer.org/).^22^

To investigate the relationship between the charge state and *m/z* range of a given peptide with its fragmentation behavior, the cross-link identifying spectra (CSMs) were sorted according to charge and *m/z* into separate groups for further comparison. Only the highest scoring CSM for each cross-linked *m/z* species (e.g. the same cross-linked peptide at different charge states) identified in the corresponding replicate was used for analysis. Sequence coverage of different groups revealed a strong dependency of fragmentation behavior on charge density, as has been previously described.^16,21^ Lower charge states (3/4+) and *m/z* (<750 for 3+ and <700 for 4+) results >70 % sequence coverage, already at low collision energies starting from 18 % NCE (Figure S2). On the contrary, a *m/z* above 700 leads to poor precursor fragmentation and therefore to a lower identification rate for low NCEs as can be seen in Figure S2 for NCE 15 & 18%. The average sequence coverage reaches a plateau between NCE 27 - 33 % and does not significantly increase with higher collision energies. When comparing the fragmentation of the two linked peptides, similar fragmentation behavior was observed. Nevertheless, the heavier/larger peptide showed a slightly better sequence coverage for all *m/z* ranges and most fragmentation energies, as it was observed for non-cleavable cross-linking reagents.^21^

The average sequence coverage as well as the scoring of all three used search algorithms indicate that most unambiguous cross-link identification can be obtained at NCEs above 27 % while the most CSMs and cross-links could be identified between 21 % and 24 % NCE.

Manual inspection of MS/MS-spectra revealed, that at least one of the two reporter ion doublets is absent at higher NCEs, while the relative intensity of the fragment ions increases with rising NCE (Figure 4 C/D). A more comprehensive investigation into the presence of reporter ion doublets confirmed these observations. While NCEs from 15 to 24 % on average yielded more than 1000 MS/MS-spectra containing at least one reporter doublet for each peptide, 39 % NCE could only generate 237. (Figure 4C) These doublets are essential for identification by the algorithm employed, which is why lower NCEs are able to identify more cross-linked peptides than higher NCEs. Unlike the number of identifications, the conversion rate of MS/MS scans containing reporter ion doublets to CSMs increases from roughly one third at 15 % NCE to 86 % for NCEs >30 %.

In the case of DSSO it has been previously assumed that CID is the most suitable fragmentation technique for reporter doublet formation.^9,15^ Therefore, HCD was compared to CID fragmentation using BSA (n=3). The previously generated inclusion list was used for this comparison, with CID (30% NCE) included as 10^th^ fragmentation event. The results show, that HCD fragmentation using lower NCEs (15-24%) results in the same number of reporter doublets as CID fragmentation. (Figure S3)

Finally, we investigated the fragmentation efficiencies of different NCEs. With energies ≥ 27 % NCE almost no precursor ion remained after fragmentation (<1 %). In contrast, the three lowest NCEs tested (15-21 %) resulted in MS/MS spectra where in average the precursor ion corresponds to 55 %, 38 % or 19 % of all detected ions, respectively. (Figure S4)

Seemingly, there is not one specific NCE that is suitable for HCD fragmentation of DSSO cross-linked peptides. However, after merging all the different results, we concluded that fragmentation energies below 21 % and above 33 % NCE can be neglected, as they do not offer unique properties or benefits but rather come with several disadvantages. Most likely, a stepped collision energy, that combines reporter ion formation (NCE 21/24%) as well as high sequence coverage (>27% NCE) in one fragmentation event will yield the optimal result, as previously described for the identification of post translationally modified peptides.^13,23,24^

Additionally, we investigated the collision energy dependent fragmentation of peptides cross-linked with the second commercially available cross-linking reagent DSBU. (for detailed procedure see Supplementary Information) On the basis of cross-links derived from BSA (n=3) the different NCEs were compared based on the number of identified cross-linked sites, their average scoring and the number of spectra containing at least one reporter doublet for both peptides. The results show, that all three parameters are significantly less influenced by the fragmentation energy compared to cross-links formed with DSSO. While DSSO cross-linked peptides are less likely to form reporter doublets when fragmented with NCEs >27% (Figure 4B), this effect is less critical for DSBU cross-linked peptides. (Figure S5) However, dramatically less cross-links are identified when too low fragmentation energies (< 21 % NCE) are used. These results indicate, that existing fragmentation strategies for DSBU already cover the optimal NCE range. Therefore, we did not further investigate the fragmentation behavior of DSBU.

### Stepped Collision Energy Comparison

Based on our observations, we tested 3 stepped collision energies that span the NCEs optimal for both doublet formation and sequence coverage (24±3 %, 27±3 % and 27±6 % NCE). For comparison a mixture of all investigated proteins was used. To allow an unbiased comparison, only precursors from an inclusion list (combined inclusion list from separate proteins see “Effect of NCE on Cross-Link identification”) were selected for fragmentation. For each precursor 3 subsequent fragmentation events were triggered.

The method using 27±6 % NCE identified the most CSMs in all three replicates. The average number of CSMs increased by more than 11 % from 24±3 % NCE to 27±6 % NCE with 27±3 % NCE identifying 6 % less. (Figure 5) Also, when using MeroX for data analysis the highest number of CSMs were identified using 27±6 % NCE. (Figure S3) To summarize, using an NCE of 27±6 % yields the highest number of identifications and therefore is the fragmentation energy of choice for MS2 based identification of DSSO cross-linked peptides.

**Figure 5.**
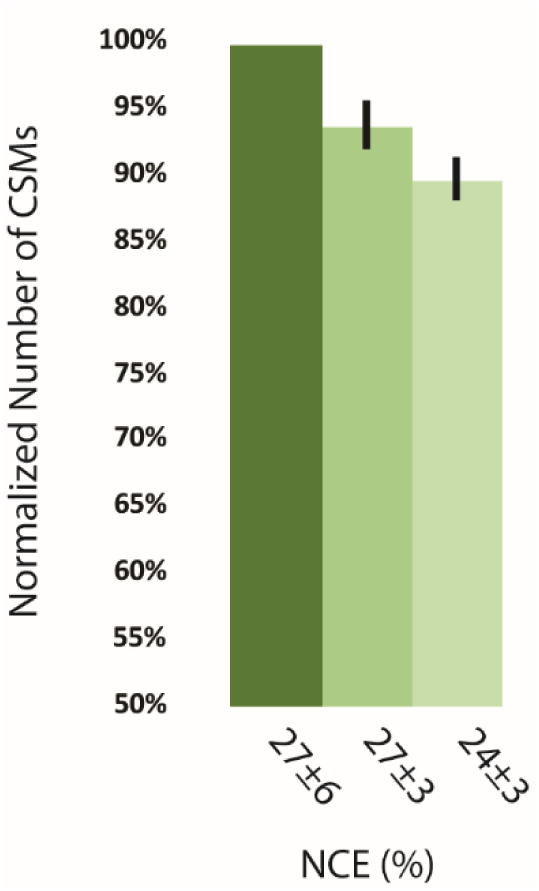
Comparison of the three most promising normalized collision energies for stepped-HCD. (n=3, Error bars represent the 0.95 CI)

### Comparison with other Methods

Stepped-HCD was compared to previously described methods using BSA and the 2.5 MDa 70S *E. coli* ribosome that represents an ideal model for large protein complexes. The cross-linked BSA digest was measured in triplicate, and for the ribosome… Since manual inspection of CSMs revealed that those with a low Δ-score (<40) mostly show poor fragment ion series, the results were filtered for a Δ-score ≥40.

In the case of BSA, the newly developed stepped-HCD method identified an average of 75 unique cross-linked sites, statistically not significantly more than MS2-MS3 that identified an average of 71. These results are in the same magnitude as suggested by a community wide XLMS study.^25^

However, both outperformed the approach employing two complementary fragmentation types CID-ETD by 56 % and 48 %, respectively. (Fig. 6A) Therefore, this approach was not included for the comparison on the more complex sample.

**Figure 6.**
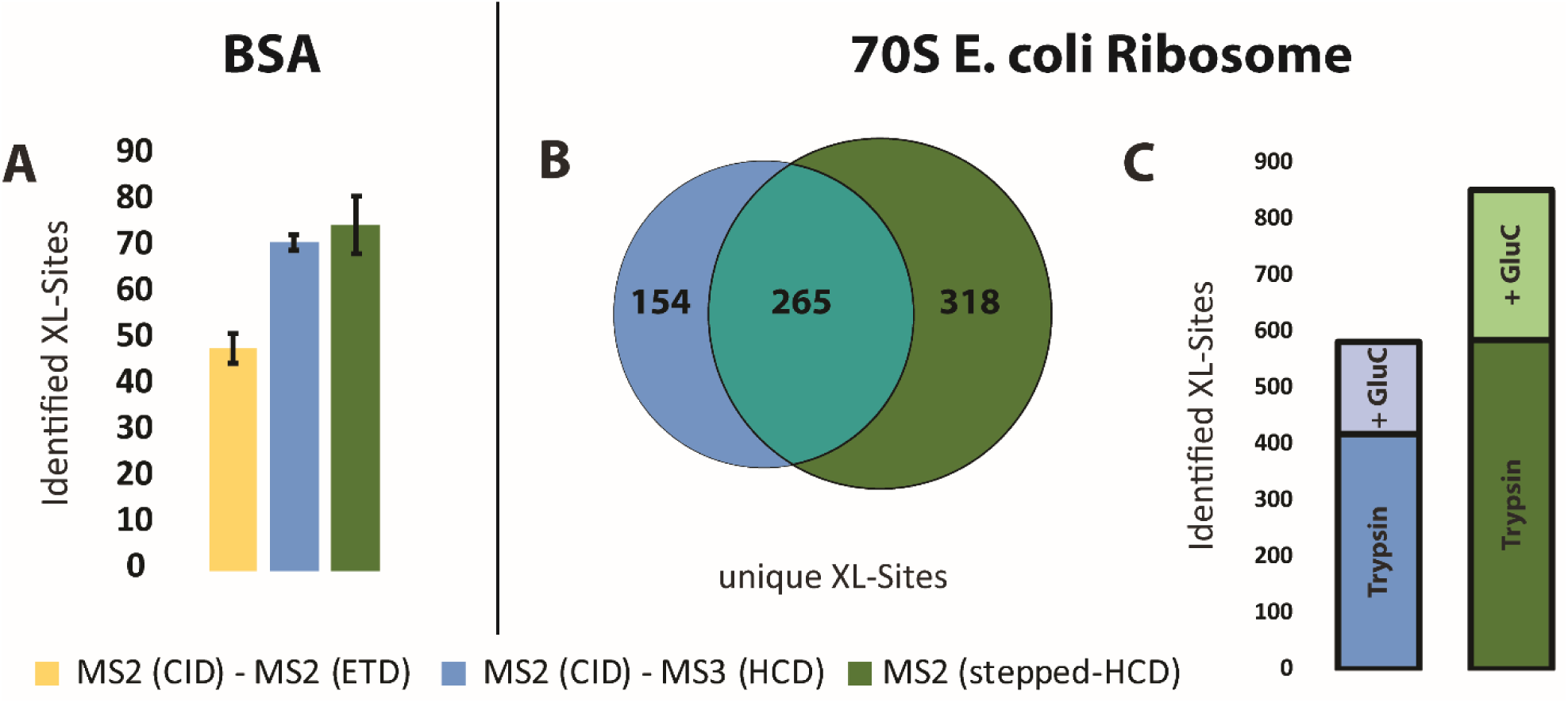
Comparison of the stepped-HCD approach with previously published methods on BSA (n=3, Error bars represent the 0.95 CI). (A) Overlap of unique XL-Sites identified in the tryptic digest using stepped-HCD and MS^2^MS^3^. (B) The additional unique XL-Sites obtained from the sequential digest using trypsin and GluC. (C)

To further evaluate the performance of the stepped-HCD it was compared to the MS^2^-MS^3^ method on cross-links derived from the 2.5 MDa 70S *E. coli* ribosome. The digest was enriched for cross-linked peptides using SEC. 583 unique cross-linked sites were identified using the stepped-HCD method applying a 1% FDR. Meanwhile, MS^2^-MS^3^ identified 28 % less (419) when using the same parameters. (Figure 6B) However, only 36 % (265) of the cross-linked sites are shared between both methods, while 318 and 154 are uniquely identified by stepped-HCD and MS^2^-MS^3^, respectively. This indicates a certain degree of complementarity of the two strategies, as was previously reported for other CID-cleavable cross-linking reagents.^26^ Hence, a combination of both strategies are likely to lead to a more comprehensive cross-linking result. The advantage of stepped-HCD over the MS2-MS3 acquisition is partially due to the inclusion of triply charged precursors; 44 unique cross-linked sites were exclusively identified using stepped-HCD and exclusively at charge state 3+. When filtering the results for triply charged precursors, stepped-HCD identified 522 unique sites and still outperforms MS^2^-MS^3^ by more than 20 %, although time was wasted acquiring 3+ precursors.

Additionally, replicate analysis of SEC-fractions 2 & 3 confirms the superiority of stepped-HCD. (Figure S7)

Recent literature highlights the benefits of sequential digestion using different proteases for more comprehensive cross-link identification through enhanced protein sequence coverage.^27^ Therefore, the DSSO cross-linked ribosome was additionally digested sequentially using trypsin and GluC. Direct comparison of the two acquisition strategies pointed out the versatility of stepped-HCD. (Figure 6C) While the MS^2^-MS^3^ method identified 195 unique cross-linked sites, stepped-HCD identified more than twice as many (431 unique sites), again at 1% FDR. This advantage is likely due to the shorter peptides produced by the sequential digest. Hence, cross-links tend to occur at lower charge states (predominantly 3+), that have not been considered for MS^2^-MS^3^. In summary, stepped-HCD allowed identification of 849 unique XL sites, 320 more than the MS^2^-MS^3^ approach.

## CONCLUSION

MS-analysis of peptides cross-linked with the cleavable cross-linker DSSO comes with several challenges.

1. The C-S bonds adjacent to the sulfoxide group are more labile than the peptide bonds, so different fragmentation energies are required for simultaneous cross-linker and peptide cleavage.
2. High fragmentation energies result in the loss of the cross-link reporter doublet ions.

Therefore, three stepped collision energies that combine higher and lower fragmentation energies were tested. The best performing acquisition strategy using 27±6 % NCE, was subsequently compared to previously described acquisition strategies. This approach was shown to be able to identify more cross-linked sites than other acquisition strategies. In the case of BSA stepped-HCD performed equally as the previously published MS^2^-MS^3^ method, while being the simpler strategy. For the 70S *E. coli* ribosome, a large multisubunit riboprotein, stepped-HCD identified 584 cross-linked sites using a tryptic digest and thereby outperformed the MS^2^-MS^3^ acquisition method that identified 417 cross-linked sites only. In addition, it proved to be compatible with sequential digests using multiple proteases, allowing a more comprehensive cross-link analysis. Altogether our novel fragmentation strategy identified almost 850 unique cross-linked sites, 45 % more than the MS^2^-MS^3^ method.

Our approach represents a powerful alternative to previously described analysis strategies for DSSO cross-linked peptides. It allows their analysis on mass spectrometers equipped with an HCD-cell without the need for ETD or MS^n^. Thereby, it will help to make XLMS available to a broader audience. Additionally, this new approach can in principle be applied to every other sulfoxide containing cross-linking reagent.^28–30^

## Supporting information

## ACKNOWLEDGMENT

This work was supported by the Austrian Science Fund (SFB F3402; P24685-B24, TRP 308-N15 and I 3686). We thank IMP and IMBA for general funding as well as all of the technicians of the protein chemistry and mass spectrometry facility for continuous laboratory support. In particular we acknowledge K. Stejskal for method development support and R. Beveridge for manuscript revision and helpful discussion.

