## Supplementary material for "Optimized fragmentation improves the identification of peptides cross-linked using MS-cleavable reagents"

### Supplementary Information

**Cross-Link Data Analysis using MeroX.**

Raw files were converted to.mgf files using Proteome Discoverer 2.2 (PD). For MeroX 1.6.6[1] the following settings were used: Cross-Linker: DSSO (+158.00376 Da, reactivity towards lysine and protein N-terminus); cross-linker fragments: alkene (+54.01056 Da), unsaturated thiol (+85.98264 Da), sulfenic acid (+103.9932 Da); alkene and unsaturated thiol were selected as essential; additionally the RISE Mode was activated to compensate for 1 missing ion; MS1 accuracy: 10 ppm; MS2 accuracy: 20 ppm; used enzyme: trypsin; max. missed cleavages: arginine 3, Lysine 3; minimum peptide length: 5; max. modifications: 2; peptide mass: 300-7000 Da; static modifications: carbamidomethylation (cysteine, +57.021 Da); dynamic modifications: oxidation (methionine, +15.995 Da). For the database search the false discovery rate (FDR) was set to 1 %.

**Investigations on DSBU cross-linked peptides**

BSA (1 mg/ml) was cross-linked with 500 µM DSBU for 30 minutes at room-temperature. The sample was further processed, as described for DSSO cross-linked proteins. Cross-links were analyzed on a QExactive HF-X mass spectrometer applying the previously published MS-method using HCD with 30±3 % NCE.[1] Digested peptides were separated using a Dionex UltiMate 3000 HPLC RSLC nanosystem prior to MS analysis. The HPLC was interfaced with the mass spectrometer via a Nanospray Flex™ ion source. For sample concentrating, washing and desalting, the peptides were trapped on an Acclaim PepMap C-18 precolumn (0.3x5mm, Thermo Fisher Scientific), using a flowrate of 25 µl/min and 100% buffer A (99.9% H2O, 0.1% TFA). The separation was performed on an Acclaim PepMap C-18 column (50 cm x 75 µm, 2 µm particles, 100 Ä pore size, Thermo Fisher Scientific) applying a flowrate of 230 nl/min. For separation, a 90 minutes solvent gradient ranging from 2-35% buffer B (80% ACN, 19.92% H2O, 0.08% TFA) was applied. Mass spectra were recorded as follows: Full scan: 60000 resolution; AGC target 1e6; 60 ms max. injection time; from 350-1500 m/z. MS/MS-spectra: 30000 resolution; AGC target 5e4; 150 ms max. injection time; from 200-2000 m/z, 3.3e4 intensity threshold. The top 15 most intense ions with charge state >2+ were selected for fragmentation and subsequently excluded for 30 seconds. For identification XLinkX 2.2 was used.[2]

Identified cross-links were put onto an inclusion list using the Proteome Discoverer 2.2 output. To investigate the NCE dependence of DSBU cross-links, the same mass spectrometer settings as described for DSSO cross-linked peptides (see Material and Methods) and the freshly generated inclusion list were used.

For XLinkX the following settings have been used: Cross-Linker: DSBU (+196.084792 Da, reactivity towards lysine); cross-linker fragments: Bu (+85.05276 Da), UrBu (+111.03203 Da), cross-link doublets: Bu/UrBu (Δ-mass 25.9793 Da); MS1 accuracy: 10 ppm; MS2 accuracy: 20 ppm; used enzyme: trypsin; max. missed cleavages: 4; minimum peptide length: 5; max. modifications: 4; peptide mass: 300-7000 Da; static modifications: carbamidomethylation (cysteine, +57.021 Da); dynamic modifications: oxidation (methionine, +15.995 Da). For the database search the false discovery rate (FDR) was set to 1%.

**Shotgun Analysis of the Ribosome Sample.**

The ribosome sample was digested as described for the cross-linked samples. The digest was separated using a Dionex UltiMate 3000 HPLC RSLC nanosystem prior to MS analysis. The HPLC was interfaced with the mass spectrometer via a Nanospray Flex™ ion source. For sample concentrating, washing and desalting, the peptides were trapped on an Acclaim PepMap C-18 precolumn (0.3x5mm, Thermo Fisher Scientific), using a flowrate of 25 µl/min and 100% buffer A (99.9% H2O, 0.1% TFA). The separation was performed on an Acclaim PepMap C-18 column (50 cm x 75 µm, 2 µm particles, 100 Ä pore size, Thermo Fisher Scientific) applying a flowrate of 230 nl/min. For separation, a solvent gradient ranging from 2-35% buffer B (80% ACN, 19.92% H2O, 0.08% TFA) over 3h was applied. Mass spectra were acquired on a QExactive HF Orbitrap-MS. Full scans were recorded at a resolution of 60000 ranging from 380-1500 m/z (AGC 1e6, max injection time 60 ms). MS/MS scans of the top 10 most intense ions with charge state 2-6+ were recorded at a resolution of 30000 ranging from 200-2000 m/z (AGC 1e5, max injection time 105 ms). Selected precursors were fragmented using HCD with an normalized collision energy (NCE) of 27 %. The Minimum AGC target was set to 5e3 with an intensity threshold of 4.8e4. Ions that have been selected for fragmentation have been excluded from MSMS for 60 seconds.

The obtained .raw file was analyzed in PD 2.1 using MS-Amanda 2.2[3] and apQuant[4] using the following settings: MS1 accuracy: 5 ppm; MS2 accuracy: 10 ppm; used enzyme: trypsin; max. missed cleavages: 2; max. modifications: 4; peptide mass: 350-5000 Da; static modifications: carbamidomethylation (cysteine, +57.021 Da); dynamic modifications: oxidation (methionine, +15.995 Da). For FDR calculation Percolator was used with a target FDR of 1%. For apQuant 5ppm mass tolerance and PSM confidence level "High" with a minimum score of 150 and a minimum peptide length of 7 was used. For protein quantification the iBAQ option was used.

For the generation of the FASTA file for cross-link search, all protein that were at least "Master Candidate" and had at least 2 identified peptides were considered.

**Supplementary Figures:**

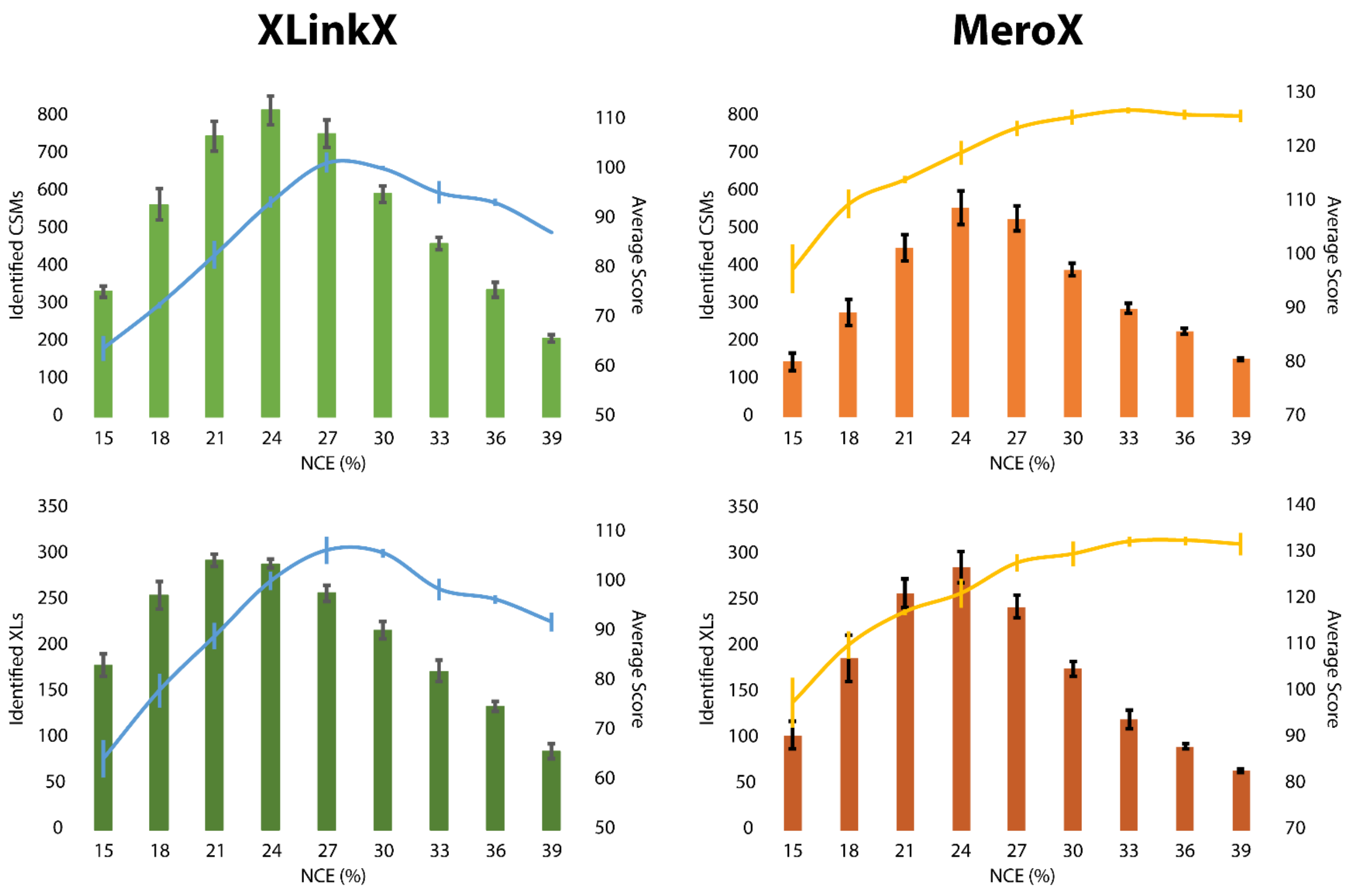


**Figure S1:** Mean number of identified cross-link spectrum matches (CSMs), unique cross-linked sites (XLs) identified with the respective NCE and their average scoring. (n=3, Error = 0.95 confidence interval [CI])

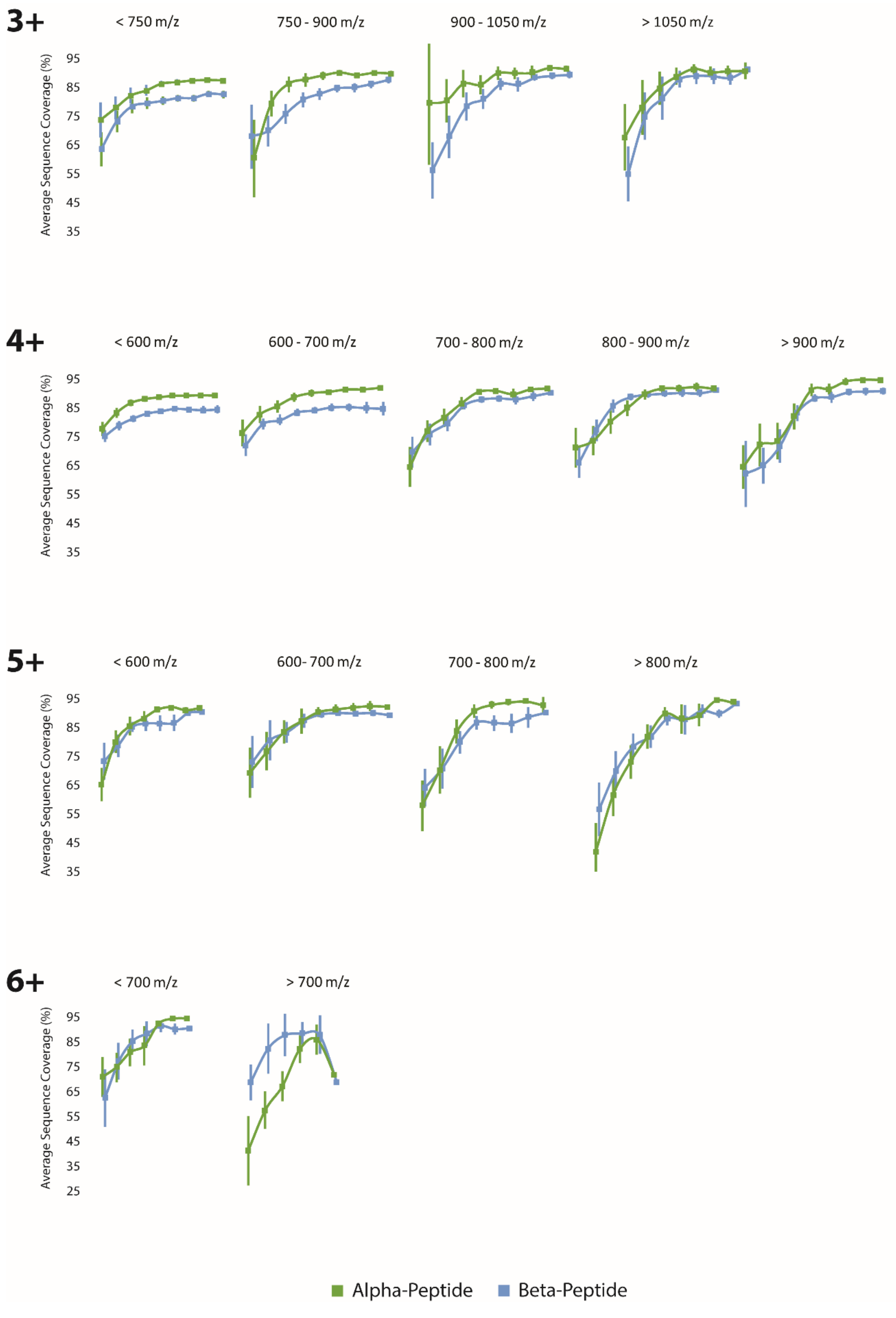


**Figure S2:** Average Sequence coverage obtained at different NCEs, for different m/z and z ranges.

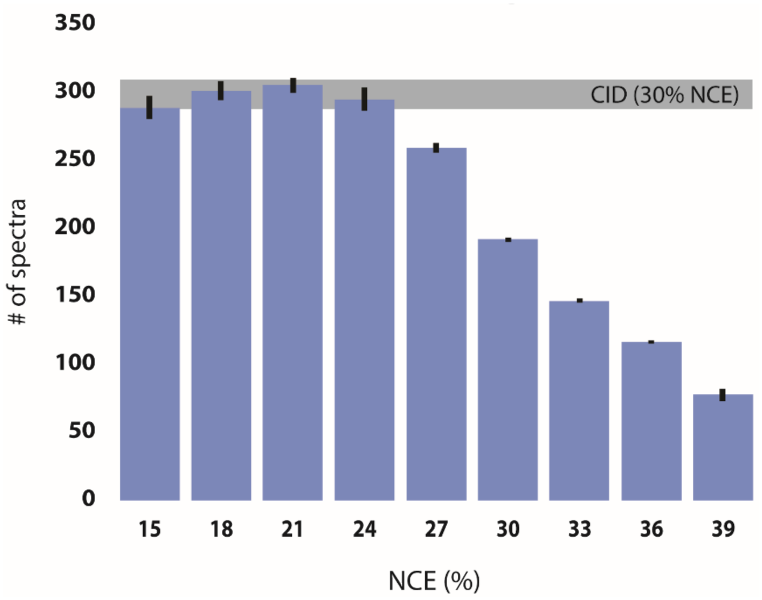


**Figure S3:** Number of reporter doublet containing spectra obtained with CID fragmentation (30% NCE) and HCD fragmentation applying NCEs ranging from 15-39%. (n=3, Error = 0.95 CI)

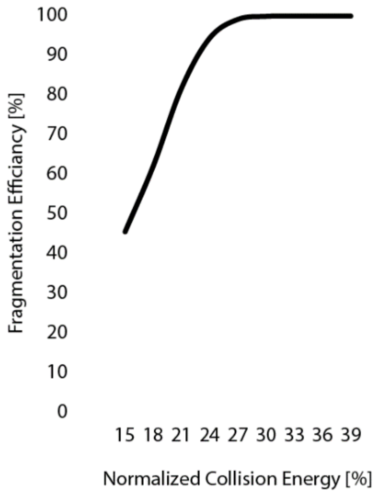


**Figure S4:** Average fragmentation-efficiency obtained at different NCEs

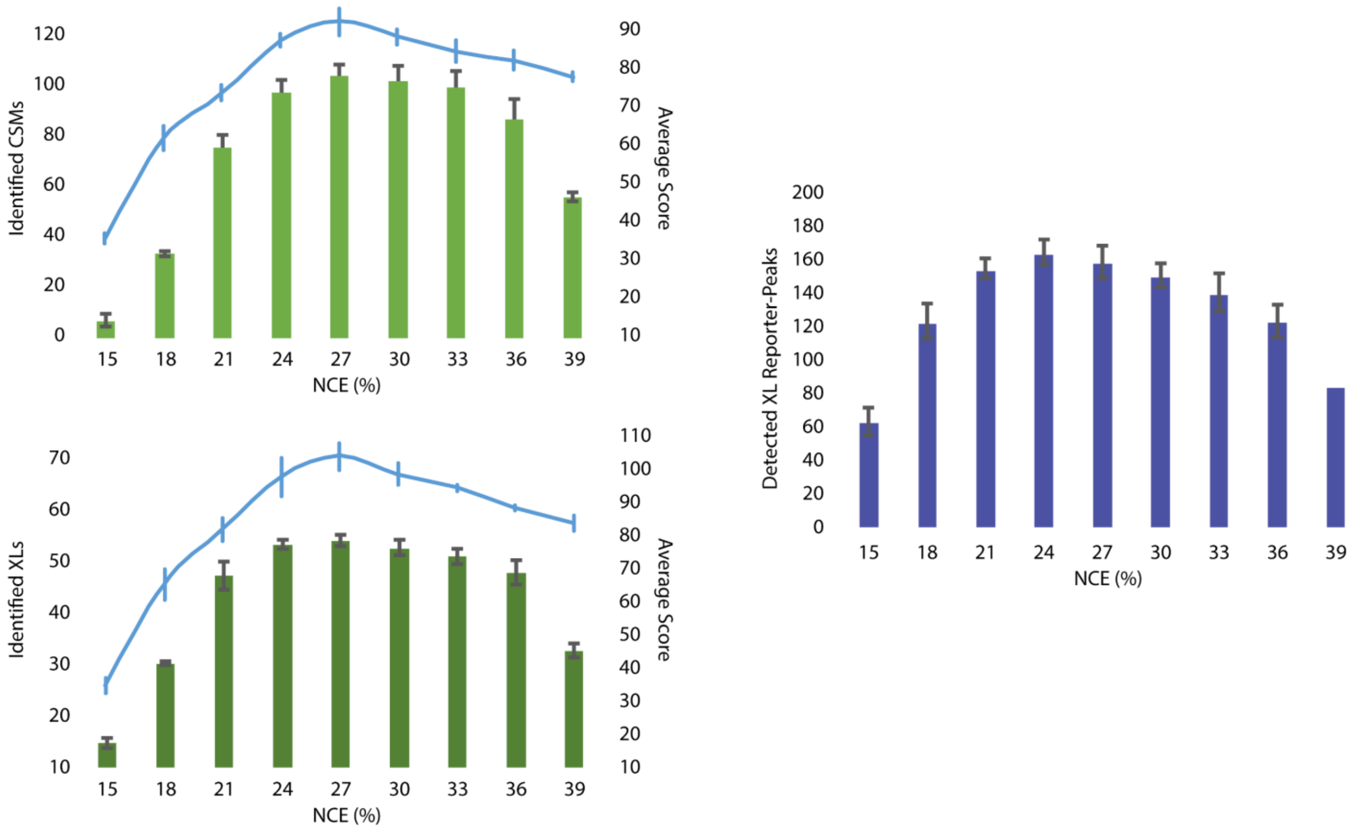


**Figure S5:** Investigation into the fragmentation energy dependent identification of disuccinimidyl dibutyric urea (DSBU) cross-linked peptides and the likelihood for reporter doublet formation. (n=3, Error = 0.95 CI). Data was analyzed using XLinkX 2.2.

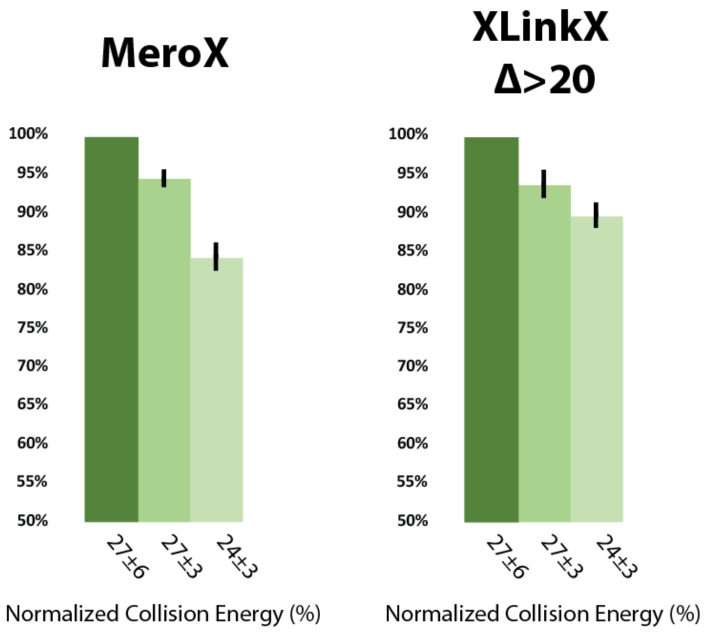


**Figure S6:** Comparison of the three most promising stepped collision energies in terms of identified spectra using the different search engines (Normalized on the number of CSMs obtained with 27±6 & NCE, n = 3, Error = 0.95 CI).

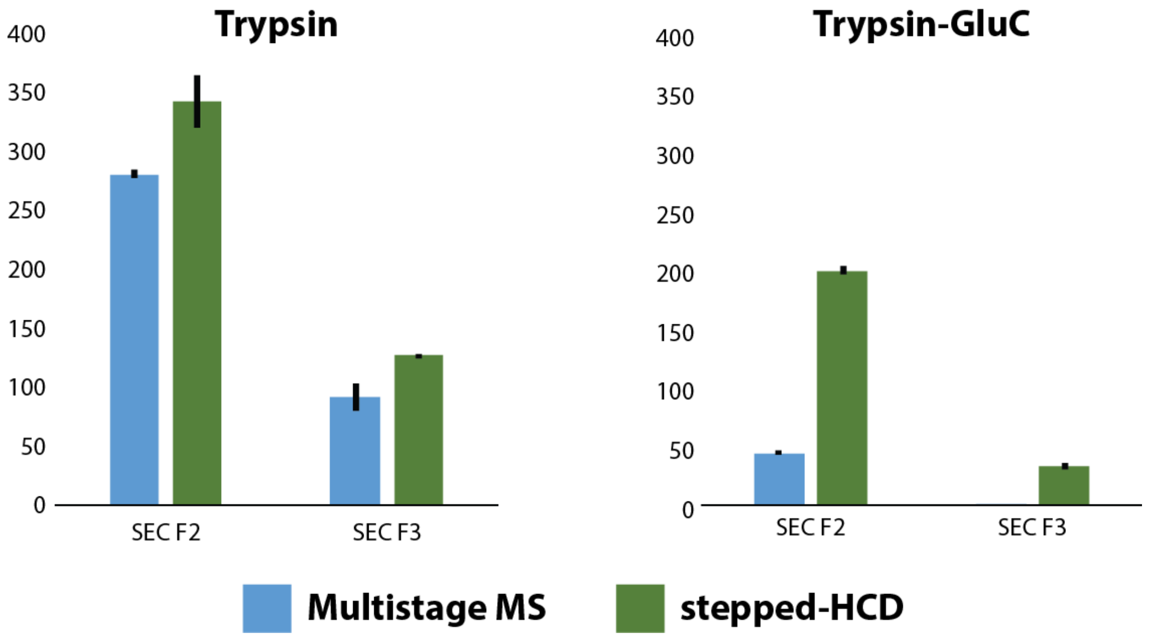


**Figure S7:** Identified cross-linked sites identified in SEC fractions 2 & 3 obtained using the different enzyme(-combinations) and fragmentation strategies, respectively. (n = 3, Error = 0.95 CI)

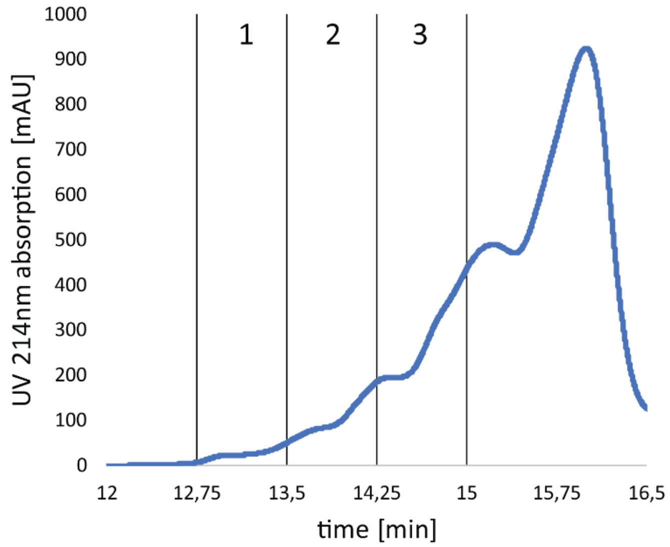


**Figure S8:** UV-chromatogram of the SEC-fractionation of the ribosomal sample indicating the three analyzed fractions.
